# Tug-of-peace: Visual Rivalry and Atypical Visual Motion Processing in MECP2 duplication Syndrome of Autism

**DOI:** 10.1101/2022.09.23.509144

**Authors:** Daria Bogatova, Stelios M. Smirnakis, Ganna Palagina

## Abstract

Extracting common patterns of neural circuit computations in the autism spectrum and confirming them as a cause of specific core traits of autism is the first step towards identifying cell- and circuit-level targets for effective clinical intervention. Studies in human subjects with autism have identified functional links and common anatomical substrates between core restricted behavioral repertoire, cognitive rigidity, and over-stability of visual percepts during visual rivalry. To be able to study these processes with single-cell precision and comprehensive neuronal population coverage, we developed the visual bi-stable perception paradigm for mice. Our task is based on plaid patterns consisting of two transparent gratings drifting at an angle of 120° relative to each other. This results in spontaneous reversals of the perception between local component motion (motion of the plaid perceived as two separate moving grating components) and integrated global pattern motion (motion of the plaid perceived as a fused moving texture). Furthermore, this robust paradigm does not depend on the explicit report of the mouse, since the direction of the optokinetic nystagmus (OKN, rapid eye movements driven by either pattern or component motion) is used to infer the dominant percept. Using this paradigm, we found that the rate of perceptual reversals between global and local motion interpretations of the stimulus is reduced in the MECP2 duplication mouse model of autism.

Moreover, the stability of local motion percepts is greatly increased in MECP2 duplication mice at the expense of global motion percepts. Thus, our model reproduces a subclass of the core features in human autism (reduced rate of visual rivalry and atypical perception of visual motion). This further offers a well-controlled approach for dissecting neuronal circuits underlying these core features.

## Introduction

Autism is a group of neurodevelopmental disorders traditionally conceptualized as impairments of high-level cognitive functions leading to deficient social communication and repetitive restricted behavioral repertoire. A distinct perceptual style accompanies these high-level features of the condition and sensory picture of the world, focusing on the fine details of the environment rather than globally integrated scenes (***Robertson and Baron-Cohen, 2017**; **Van der Hallen et al., 2019***). Even before social deficits become evident, over 90 % of individuals with autism experience altered sensation and atypical sensory perception that affect every sensory modality (***Grzadzinski et al., 2013**; **Robertson and Baron-Cohen, 2017**; **Simmons et al., 2009**; **Van der Hallen et al., 2019**;**Robertson and Simmons, 2015)***. A recent version of DSM (2013) now lists atypical sensory perception as a core diagnostic feature of autism, together with social communication deficits and restricted repetitive behaviors (***APA, 2013***). Another relevant feature of autism is the heterogeneity of expression of core traits and remarkable behavioral diversity across individuals, affecting all aspects of interaction with physical and social environments (***Baron-Cohen et al., 2009**; **Bolton et al., 2020**;**Lawson et al., 2015**; **Robertson and Baron-Cohen, 2017**; **Shafritz et al., 2008**; **Uddin, 2021**;**Van der Hallen et al., 2019)***. Thus, to explain the autistic brain, one must consider what it is in the brain that provides common ground for apparently disparate phenomena such as social communication, cognitive rigidity, and atypical visual perception. What can account for the phenotypical diversity of the condition and, at the same time, ensure the presence of its core features in most affected individuals? Importantly, it becomes critical to develop behavioral paradigms and approaches that can reliably measure these putative common ground processes and be applied in mouse models of autism with the long-term goal of studying the circuit basis of the condition and providing a pipeline for fast drug candidate screening.

In this work we apply a bi-stable visual perception paradigm to study the mouse model of MECP2 duplication syndrome (***Collins et al., 2004**; **Ramocki et al., 2010***), a syndromic ASD caused by genomic duplication of methyl-CpG-binding protein 2 (***Ramocki et al., 2010***) that exhibits 100% penetrance in males. In humans, MECP2 duplication syndrome displays all core features of idiopathic autism (***Peters et al., 2013**; **Ta et al., 2022***). MECP2 duplication mice carry a number of core autism features including repetitive stereotyped behaviors, altered vocalizations, increased anxiety, motor savant phenotype and over-reliable visual responses (***Collins et al., 2004**; **Jiang et al., 2013**; **Samaco et al., 2012**; **Sztainberg et al., 2015**; **Zhang et al., 2017**; **Zhou et al., 2019**;**Ash et al., 2017**, **2021b**,a, **2022***).

Bistable visual perception paradigms are a natural choice for studying autistic brains. First, the dynamics of visual rivalry are altered in idiopathic human autism, with subjects showing a decreased rate of perceptual reversals (***Robertson et al., 2013**; **Spiegel et al., 2019***). Second, visual rivalry is a distributed computation involving both low-level sensory cortical areas and high-level association areas, such as the secondary motor cortex and prefrontal cortex ***(Kleinschmidt et al., 1998**; **Knapen et al., 2011**; **Leopold and Logothetis, 1996**,**1999**; **Lumer et al., 1998**;**Lumer and Rees, 1999)***. Thus, its dynamics are based on stimulus representation sub-networks in the early visual cortex as well as visuomotor areas and high-level cognition-related non-sensory sub-networks of higher-order cortical areas. As a result, it can be a suitable candidate method to evaluate both (a) low-level sensory processing dysfunction that involves the primary sensory cortex, and (b) high-level dysfunction such as cognitive rigidity and restricted social communication, which rely on distributed computations in non-sensory association frontal and prefrontal cortical areas. Indeed, in human autism slower rate of bistable alternations was shown to share an anatomical substrate with general cognitive rigidity, and binocular rivalry phenotype predicts the severity of social phenotype and the diagnosis of ASD (***Spiegel et al., 2019**; **Watanabe et al., 2019***). Importantly, it was suggested that the dynamics of the visual rivalry are dependent on brain-wide excitatory-inhibitory balance — a process that is also proposed to be altered in autism, leading to the expression of core traits of ASD (reviewed in (***Zhao et al., 2021***)). Finally, our visual rivalry paradigm utilizes a bistable moving plaid, in which the subject’s perception switches between the local motion-based, “transparent” interpretation of the stimulus versus the global motion-based, “coherent” interpretation. Thus, our paradigm also offers the additional advantage of exploring another core trait of autistic brains: atypical processing of visual motion and detail-oriented sensory processing style (reviewed in (***Robertson and Baron-Cohen, 2017**; **Van der Hallen et al., 2019***)).

Using our paradigm, we found that MECP2 duplication mice recapitulate the phenotype in a subset of subjects with idiopathic autism. Specifically, compared to unaffected littermates, MECP2 duplication mice display a reduced rate of perceptual reversals during visual rivalry and strongly prefer to focus on local moving cues rather than the integrated percept of coherent global motion.

## Experimental Procedures

### Animals

All experiments and animal procedures were performed in accordance with guidelines of the National Institutes of Health for the care and use of laboratory animals and were approved by the Brigham and Women’s Hospital (BWH) Institution Animal Care and Use Committee (IACUC). We used mice of two different backgrounds: mixed background C57×FVB-MECP2 duplication mice and 129-MECP2 duplication mice (***Ash et al., 2022***). Mixed background mice were produced by crossing C57Bl6J mice to FVB-MECP2 duplication line (Tg1) (***Collins et al., 2004***) mice to generate F1 C57×FVB-MECP2 duplication mice and non-transgenic littermate controls. Experiments were performed in 4–6-month-old animals. Cohorts were balanced in terms of animal sex (129 background: 3 male and 3 female pairs; C57×FVB background: 4 male and 4 female pairs). The experimenters were blind to animal genotypes during experiments and analysis.

### Surgery

All procedures were performed according to animal welfare guidelines authorized by the Brigham and Women’s Hospital IACUC committee. Micewere anesthetized with 1.5 % isoflurane. The mouse head was fixed in a stereotactic stage (Kopf Instruments), and eyes were protected with a thin layer of artificial tears ointment (GenTeal). The scalp was shaved and disinfected by applying consecutive swabs of the povidone-iodine solution and 70 % ethanol, and then the scalp was resected. A custom-made titanium head plate was attached to the skull with dental acrylic (Lang Dental), preventing occlusion of the mouse’s visual field.

### Visual stimulation

Visual stimuli were generated in MATLAB and displayed using Psychtoolbox (***Brainard, 1997***). The stimuli were presented on two LCD monitors with a 60 Hz frame rate, positioned ≈ 10 cm in front of the right eye and covering 180° of the right visual field of the mouse. The screens were gamma-corrected, and the mean luminance level was photopic at 80 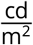. Visual stimuli consisted of drifting square-wave gratings and plaids of 120° cross angle composed of the grating stimuli components. The gratings had the following parameters: temporal frequency 1.7 Hz, spatial frequency 0.06cycles/°, spatial duty cycle 0.8 (white bar set to 60%, black bar set to 40%). These parameters were selected to accommodate average spatial frequency and velocity preferences in visually-responsive neurons across mouse visual cortical hierarchy (***de Vries et al., 2019**; **Gao et al., 2010**; **Niell and Stryker, 2008**; **Ohki et al., 2005***). Additive plaid patterns were constructed by summing up component gratings of 50° contrast (***Smith et al., 2005***). Each instance of plaid or grating movie was preceded by a gray isoluminant screen for 5 min. We kept mean luminance constant throughout both the background and the stimulation periods.

### Optokinetic nystagmus

We recorded optokinetic eye movements (EM) elicited during observation of drifting gratings and plaids in 13 head-posted mice MECP2 duplication — littermate pairs. Seven pairs were C57×FVB mixed background mice and six pairs were 129 background mice. The stimulus was presented on two screens covering ≈ 180° of the visual field of the mouse. The center of each screen was located at 10 cm from the mouse (Figure 2). We used an infrared camera (model MAKO U-29, Allied Vision Technologies) and a hot mirror to record the movements of the right eye at 300 Hz. We analyzed 5 to 15-min-long movies off-line with Deep Lab Cut toolbox (***Mathis et al., 2018***) to detect the pupil and extract its diameter and position. Optokinetic eye movement is composed of smooth pursuit following the motion of salient features in the stimulus, followed by a rapid saccade in the direction opposite to the direction of the global stimulus drift to stabilize the image on the retina (***Cahill and Nathans, 2008***). This pattern of movements (slow pursuit phase plus rapid saccade phase) repeats as long as the stimulus (drifting grating or plaid) is present and is attended by the animal. We analyzed both vertical and horizontal EM components to classify plaid-induced OKN as aligned with local motion percept vs. aligned with global motion percept. Periods containing eye-blink artifacts and mouse grooming, that the Deep Lab Cut algorithm identified as having a probability of being a pupil below 95%, were removed from the analysis. We applied a linear fit to the slow pursuit phase of each EM and calculated the eye movement amplitude from each fit (Figure 1C). The direction of each EM was determined by comparing the amplitude of horizontal and vertical saccade projections of EM components. We then plotted histograms of the directions of EMs around 0°, which corresponds to the horizontal direction (the direction of the drift of the global stimulus; see Figure 1). For the plaid-induced OKN, we classified each EM as component- or pattern-aligned. For this, we first determined the horizontal direction and the average width of the distribution of EM angles evoked by horizontally drifting gratings. We used one standard deviation (SD) from the mean as the threshold for pattern-aligned EM angles. Thus, any EM whose angle exceeded this threshold was classified as component-aligned, while EMs with angles inside the [-SD, +SD] interval are classified as pattern motion-aligned (Figures 1E and 1G).To study the dynamics of OKN alternations between following the global pattern motion or following component motion, we analyzed 2-615 min movies of OKN induced by the plaid stimulus moving in the temporonasal (T→ N) direction. We extracted the periods of the stable OKN (at least two saccade-pursuit pairs, occurring without a break between the pairs; e.g., saccadic movement is followed by the pursuit phase of the next pair). To identify periods without the OKN (breaks), we first examined the distribution of lengths of pursuits of individual nystagmoid eye movements. Periods of eye drift without return saccades exceeding the 95^th^ percentile of this distribution of lengths were considered breaks in the OKN. During breaks mouse either was not attending to the stimulus and thus experienced no OKN, closed eyes, or experienced eye blinks and grooming bouts. Periods of OKN between breaks (OKN epochs) had to contain at least two consecutive saccade-pursuit pairs to be accepted for the analysis of perceptual reversals. Each movie had to contain at least 3 min of OKN to be accepted for the analysis.

**Figure 1:**
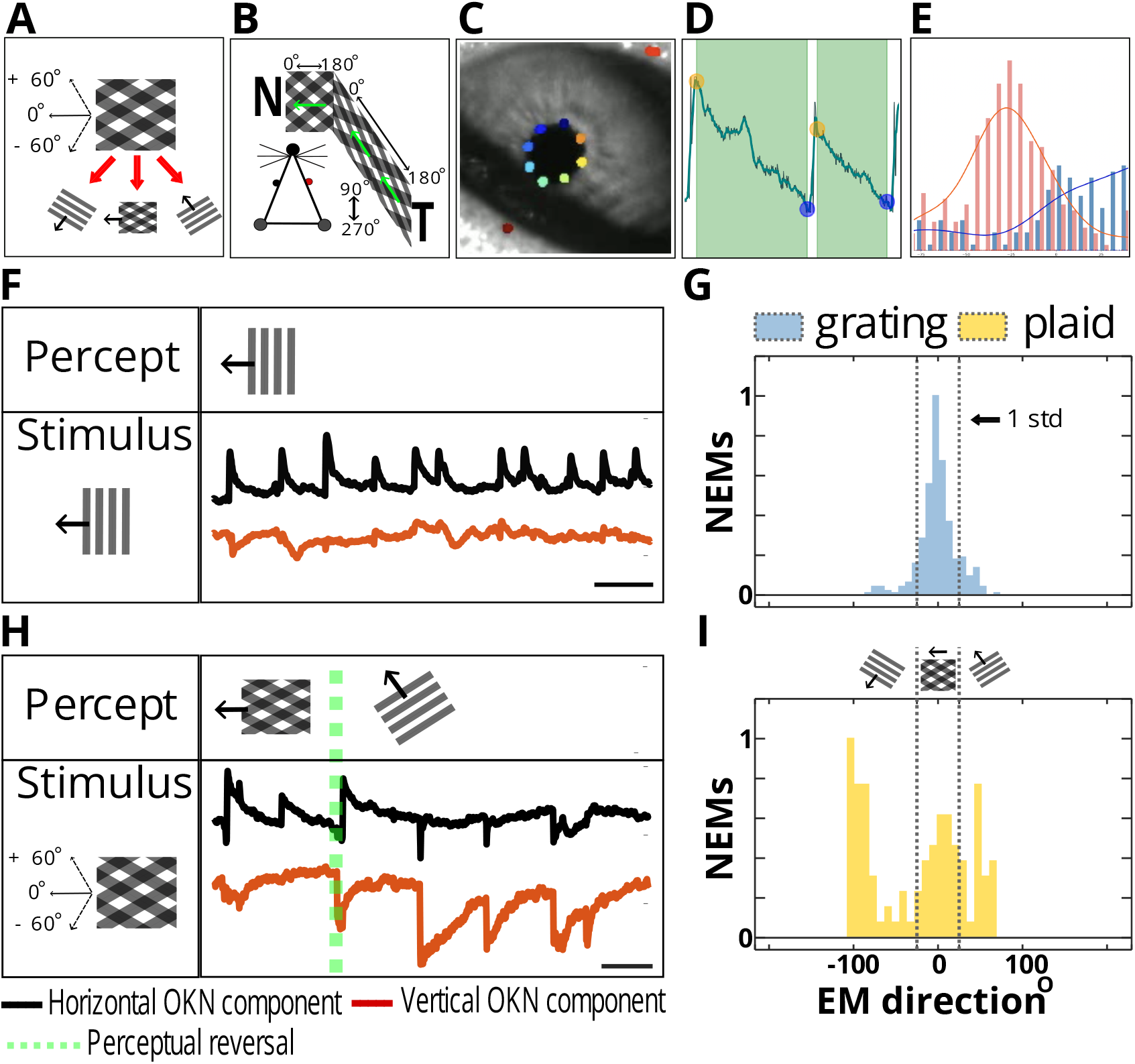
Bistable OKN responses under visual rivalry. A. Bistable moving plaid stimulus. Type I symmetric plaid is composed by summing two 50 % contrast component gratings. The gratings move at an angle of 120 degrees relative to each other. This plaid can be seen either as two individual gratings moving at an angle or as a sum of gratings integrated percept of pattern motion. The direction of pattern motion lies in between the directions of motion of each grating. Thus, the observer can follow three directions of motion (lower panel): pattern motion (direction set at 0°), and either of the component grating’s drift, offset at +60° and −60° from the vector of the plaid’s motion (insets). **B. Experimental setup.** We presented the stimuli on two screens positioned at equal distances from the mouse head to cover 180° of the mouse ipsilateral visual field. We head-posted the mouse to prevent head movements and monitored eye movements with an infrared camera. The mouse could walk freely on the free-moving wheel. Green arrows indicate the direction of the global drift of the stimulus. The stimulus was moving toward the mouse’s nose to induce robust optokinetic movements. **C, D, E. Data preprocessing pipeline. C**: An infrared image of the mouse eye. OKN images were collected at 300 Hz and 20 randomly selected mouse pupil movies were used to train Deep Lab Cut ResNET-150 model to extract the position and size of the animal’s pupil during the OKN. (Colored dots DeepLabCut feature detection). **D**: The vertical and horizontal components of the OKN were sorted into saccade-pursuit eye movement pairs, and eye-blink and grooming-related artifacts were located using custom-written Python toolbox “Dolia” and excluded from analysis (***Bogatova et al., 2023***)‘ manuscript in preparation. **E**: The pursuit phases of the OKN eye movements were fitted with a linear polynomial fit. The example of the ratio between fitted vertical and horizontal component of each eye movement was then used to determine its direction (angle). Pink and blue histogram show the example distributions of eye movement directions from two different 15 minute OKN movies.

**Figure 2:**
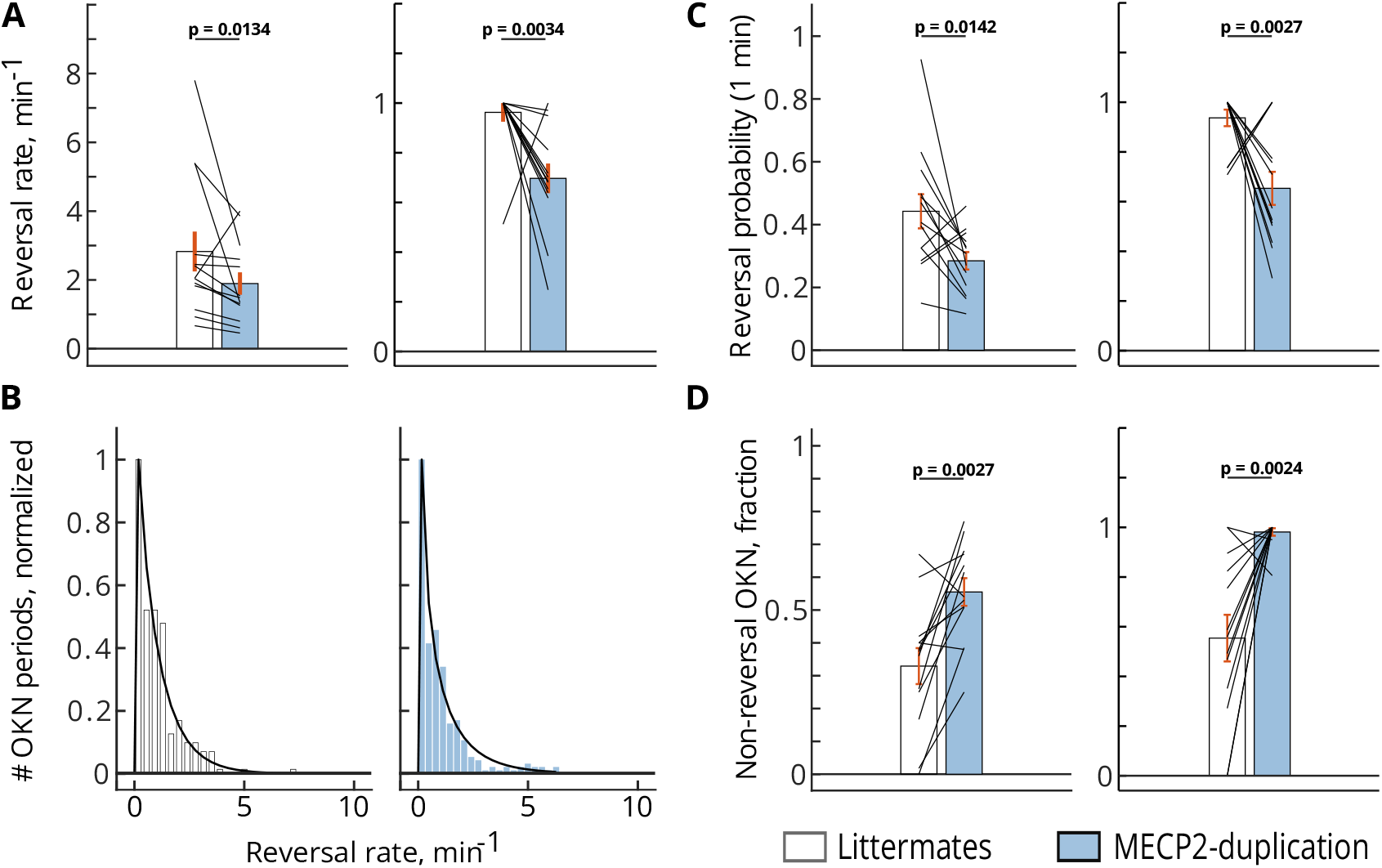
MECP2 duplication syndrome results in reduced perceptual reversal rate during visual rivalry. white bars — littermates; blue bars — MECP2 duplication syndrome. **A. The reversal rate (per minute of OKN) is consistently lower in MECP2-ds than in normal littermates.** Left panel — raw data, right panel — data normalized by maximum inside each littermate — MECP2 duplication pair. Reversals per minute: littermates, mean±sem: 2.8±0.58, median: 2.05; MECP2-ds, mean ± sem: 1.895 ± 0.325, median: 1.485. **B. The distribution of perceptual reversal rates of individual OKN periods.** Left panel (white bars) — littermates; right panel (blue bars) — MECP2 duplication. The distributions follow gamma distribution fit (littermates: *p* < 0.0001; MECP2-ds: *p* < 0.0001, *χ*^2^ test). Data were pooled across OKN periods belonging to 13 littermates and 13 MECP2 duplication animals, respectively. Before pooling, each animal’s dataset was normalized by its mean rate. **C. In accordance with the reduced reversal rate in MECP2 duplication, the probability of observing a switch after 1 minute of ongoing plaid-induced OKN was also reduced in MECP2 duplication mice.** The left panel indicates raw data, while the right panel shows the data normalized by maximum inside each littermate — MECP2 duplication pair. Reversal probability: littermates, mean±sem: 0.4425±0.054, median: 0.407; MECP2-ds,mean±sem: 0.284±0.028, median: 0.308. **D. MECP2 duplication mice consistently show a substantial fraction of OKN periods where no reversals occur, and the animal persistently tracks either pattern (“coherent” percept) or component direction (“transparent” percept).** Left panel — raw data, right panel — data normalized by maximum inside each littermate — MECP2 duplication pair. Non-reversal OKN fraction: littermates, mean±sem: 0.33±0.055, median: 0.374; MECP2-ds, mean±sem: 0.555±0.04, median: 0.54. All *p*-values are determined by two-sided Wilcoxon signed rank test (WSR), *n* = 13 pairs.

We determined the following parameters:

1. Perceptual reversal rate in each OKN epoch. The rates were averaged over epochs and movies to obtain a median value per animal.
2. The probability of experiencing a switch within 1 min of the beginning of bistable OKN.
3. The durations of “coherent” and “transparent” OKN periods in each animal. Durations were averaged over movies and animals to obtain one median value per animal.
4. Fraction of eye movements aligned with pattern and component motion across all movies of a specific animal.
5. Fraction of OKN epochs with no observed perceptual reversals (non-reversal OKN epochs).

### Statistical tests

Comparing the per-animal reversal rates, dominance period durations, and component/pattern motion ratio we used paired Wilcoxon signed rank (WSR) test comparing the 2-duplication mouse to his littermate. Statistics were computed across animals. The distributions of switch rates per OKN epoch and durations of dominance periods were fitted with the gamma distribution function. To accept or reject the fit for the gamma distribution fitting of dominance duration periods and switch rates, we used the *χ*^2^ test.

## Results

### Report-free bi-stable perception paradigm

A reliable way to infer the perceptual state when a bistable visual motion-based stimulus is presented is to measure the direction of the optokinetic nystagmus elicited by the different directions of drift generated by the rivaling stimuli (***Enoksson, 1963**; **Fox et al., 1975**; **Leopold et al., 1995**;**Naber et al., 2011**; **Watanabe, 1999**; **Wei and Sun, 1998**; **Logothetis and Schall, 1989***). Unambiguous fully coherent full-field moving visual stimuli, such as dot fields, coherently moving natural scenes and high-contrast drifting gratings, induce optokinetic nystagmus (OKN) reflex in vertebrate animals such as mammals, birds and fish (***Cahill and Nathans, 2008***). The OKN is required for the stabilization of retinal input under the conditions of a drifting visual environment. OKN eye movements consist of a slow pursuit in the direction of the stimulus followed by a fast saccade returning the eye to its initial position. OKN has been extensively validated as a reliable indicator of the dominant percept in experimental designs involving ambiguous stimuli, such as binocular rivalry ***(Fox et al., 1975**; **Naber et al., 2011**; **Watanabe, 1999**; **Wei and Sun, 1998**; **Logothetis and Schall, 1989***). Under ambiguous visual conditions, the direction of pursuit during slow phases of OKN is aligned with the direction of motion of the dominant percept (***Palagina et al., 2017***).

We have previously shown that mice can exhibit visual bistable perception when exposed to a moving transparent additive plaid stimulus covering ≈ 270° of the visual field ***(Palagina et al., 2017)***. The symmetric transparent additive plaid we used is composed of two transparent gratings of equal contrast and velocity moving at an angle to each other. Under the range of bi-stability promoting stimulus properties, the subjective perception of this stimulus alternates between the “transparent” interpretation, where two full-field component gratings slide on top of each other, and the “coherent” interpretation, where a fused pattern drifting in a direction half-way between the directions of component gratings is seen (***Adelson and Movshon, 1982**;**Moreno-Bote et al., 2010***). Large cross-angle between the grating components of the plaid, “transparency-promoting” intersection luminance values of the dark bars (equal to the sum of the luminances of the components), high component grating velocity, asymmetric intersections (occurring when the cross angle between component gratings is above or below 90°) promote a transparent interpretation (***Moreno-Bote et al., 2010**; **Movshon et al., 1985***). Symmetry in component gratings, contrast, spatial frequency and velocity favor the coherent percept (***Adelson and Movshon, 1982**;**Yo and Demer, 1992)***. In the previous work, using stimuli fulfilling these criteria (60° or 120° cross-angle between components, contrast normalization, drift velocity 2 cycle/° of visual field, spatial frequency 0.05 cycle/° and symmetry in the properties of component gratings) we were able to elicit bistable OKN in C57 wild-type mice. These properties were tailored to be optimal for mouse area V1 ***(Gao et al., 2010**; **Niell and Stryker, 2008**; **Ohki et al., 2005***). In the present study we modified the stimulus keeping in mind the necessity to drive as a large proportion of neurons as possible in different visual areas, which have varying preferences for the drift velocity and spatial frequency of the stimuli. To do this, we changed the duty cycle of the stimuli to 0.8 cycle/° and drift velocity to 1.7 cycle/°, while keeping the components symmetric (spatial frequency 0.06 cycle/°) and contrast normalized to achieve transparency-promoting luminance of the intersections. We used 120° CA (cross-angle) plaids as this was shown to induce an equidominant state (where the observer spends nearly equal time on transparent and coherent percepts) in both human observers (***Moreno-Bote et al., 2010***) and mice (***Palagina et al., 2017***). We also reduced the coverage of the visual field to 180° of the right eye’s visual field, as this was shown to induce reliable OKN in mice (***Cahill and Nathans, 2008***) while allowing us to combine the behavioral task with 2-photon imaging or electrophysiological recordings in future experimental work.

Figure 1: Using the ratio of the vertical and horizontal components, amplitudes, the directions of the eye movements were determined (see Methods section for details). Right: distributions of the directions of pursuit phases of OKN for two different OKN movies. **F, G. Grating-induced OKN.** To determine the location of zero direction (pattern motion direction) and classify eye movements as aligned with pattern motion or alternatively the motion of the components, we used OKN data obtained by presenting the 0 direction grating moving in temporonasal direction, similarly to the plaid setup. Since such a grating has only one unambiguous direction of drift, it is possible to use the mean of the eye movement direction distribution as a zero direction. Additionally, [−standard deviation, +standard deviation] can be set as a bracket in which most eye movements aligned with zero direction fall Figure 1E. The grating-induced OKN is shown in **F**: as expected, OKN eye movements contain a sole horizontal component (black trace), with no consistent vertical deflections (red trace), and this stimulus does not result in visual rivalry as only one interpretation of the stimulus is possible. In **G**, the distribution of grating-induced OKN is shown (yellow histogram, 13 zero-direction grating movies from 13 animals were used to determine zero position, and the standard deviation bracket for eye movement classification). [-standard deviation, +standard deviation] interval around the zero direction is then applied to plaid OKN data: the eye movements with directions inside this interval are classified as pattern-motion aligned, while eye movements with directions outside of this interval are classified as component-motion aligned Figure 1G, blue histogram. **H, I. Plaid-induced OKN shows bi-stable reversals of the eye movement directions.** In **H**, the mouse can follow either the plaid or the grating direction while observing the unchanging plaid stimulus. Green dotted line — location of the perceptual switch, defined as the start of the saccade where the animal starts following a different stimulus interpretation. Initially, the animal follows a pattern motion direction, as evident from OKN parameters — the presence of a robust horizontal component and no consistent vertical component (before the green dotted line). After the reversal, a solid vertical component appears (red trace) as the animal stops following the pattern motion and starts following the +60° degrees component. **I**, blue histogram: the EM directions distribution of OKN induced by a plaid stimulus. Gray dotted lines correspond to the pattern-component OKN classification bracket derived from grating OKN data (see the blue histogram in **G**). In contrast to grating data, plaid OKN, in addition to the central peak corresponding to pattern-motion aligned eye movements, has two additional peaks located at approximately +60° and −60° off the central peak and corresponding to component motion-aligned OKN.

Under the updated conditions we show that both 129-background and C57×FVB mixed back-ground mice show bi-stable optokinetic nystagmus, aligned either with the direction of component gratings or the direction of coherent pattern motion (Figure 1), similarly to what we observed in C57 mice previously (***Palagina et al., 2017***). We observed no difference in the rate of generation of OKN between littermates and MECP2 duplication mice or in the magnitude of eye movements (eye movement amplitude, arbitrary units: littermates, 7.43 ± 0.5, MECP2-ds, 6.33 ± 0.51; *p* = 0.308, WSR; OKN rate 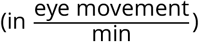: littermates, 9.7±1.7, MECP2-ds, 7±0.75, *p* = 0.216, WSR). There was no difference between 129 background animals and C57×FVB background animals in terms of OKN properties and dynamics of visual rivalry, thus these two groups were pooled together. The experimental setup is shown in Figure 1. Stimuli were presented on two contiguous screens covering 180° of the mouse contralateral visual field, and pupil position was recorded with the help of hot mirror and an infrared camera. Figure 1F shows an example of OKN elicited by a vertically oriented grating moving from the temporal to nasal direction. In this case the eye movements elicited by the stimulus are aligned with the horizontal direction (0°, taken along the temporal→nasal direction). In contrast to the unambiguous horizontally drifting gratings, OKN eye movements elicited by a 120° CA plaid show a tri-modal distribution of eye movement directions: a considerable fraction of eye movements is aligned with one of the two component grating directions in addition to the horizontally aligned OKN that corresponds to the fused pattern motion percept (Figure 1H). This strongly suggests that the perception of the mouse alternates between pattern and component motion for our stimuli in the recorded cohort of mice.

### MECP2 duplication mice show reduced rate and probability of perceptual reversals

We next examined the dynamics of bi-stable reversals between “coherent” interpretation OKN (mouse tracking global pattern; OKN eye movements aligned with the global pattern direction) and “transparent” interpretation OKN (mouse tracking the component gratings; OKN eye movements aligned with the direction of drift of either component grating) in MECP2 duplication animals ver-sus unaffected littermates. Both MECP2 duplication animals and littermates displayed bi-stable reversals. However, in MECP2 duplication syndrome mice the rate of reversals was reduced compared to their normal littermate pairs (Figures 2A to 2C), and MECP2 duplication animals displayed more frequent OKN epochs where only a single interpretation of the stimulus was consistently followed and no perceptual reversals occurred (Figure 2D; non-reversal OKN fraction: littermates, mean ± sem: 0.33 ± 0.055, median: 0.374; MECP2-ds, mean ± sem: 0.555 ± 0.04, median: 0.54; *p* = 0.0027, WRS). Consequently, the fraction of OKN epochs showing bi-stable reversals was reduced in MECP2 duplication animals. Littermates showed on average 2.8 reversals per one minute of OKN movie (mean ± sem: 2.8 ± 0.58, median: 2.05), while MECP2-ds mice showed 1.9 reversals per minute (mean ± sem: 1.895 ±0.325, median: 1.485), a significant reduction in bi-stable reversal rate (*p* = 0.0134, WSR, *n* = 13 pairs) (Figures 2A and 2B). The probability to observe a switch after 1 minute of uninterrupted plaid-driven OKN was consequently reduced in MECP2 duplication mice (littermates, mean±sem: 0.4425±0.054, median: 0.407; MECP2 duplication mean±sem: 0.284±0.028, median: 0.308; *p* = 0.0142, WSR) (Figure 2C). In sum, the properties of bi-stable reversal dynamics are altered in MECP2 duplication mice, with duplication animals showing increased proportion of reversal-free OKN epochs and reduced reversal rate and probability.

### Local versus global motion processing in MECP2 duplication mice and increased stability of local motion “transparent” percepts

The slower rate of rivalry in MECP2 duplication mice was accompanied by pronounced changes in the processing of visual motion. Specifically, MECP2 duplication animals showed strong preference for the component motion compared to their normal littermates (Figure 3). The latter either spent approximately equal time following component gratings vs. coherent pattern direction, or showed preference for coherent pattern direction. This effect was seen both in the total fraction of OKN eye movements aligned with the component versus the pattern directions (Figure 3), and in the duration of component-versus pattern-dominance periods (Figures 3B and 3C). Interestingly, for dominance periods, the strongest effect was observed in the duration of component percepts, which were on average twice as long in MECP2 duplication animals as in littermate controls (Figure 3C, littermates, mean±sem: 27.5±7.11, median: 20.2; MECP2-ds, mean±sem: 49.2±11.5, median: 31). This effect was highly reproducible across pairs and highly significant (Figure 3C, *p* = 0.0081, WSR). In contrast to the component-aligned OKN periods, the durations of pattern motion-aligned OKN periods showed disparate effects: in some duplication-littermate pairs MECP2 duplication led to an increase in pattern percept durations, while in other a decrease was observed (Figure 3D). Pooled data, including both durations of component and pattern motion-aligned OKN showed a net increase in dominance period durations, consistent with a reduced rate of perceptual reversals (Figure 3B, littermates, mean±sem: 25.2±4.5, median: 21; MECP2-ds, mean±sem: 36.7±8, median: 25; *p* = 0.0342, WSR). In addition, MECP2 duplication animals showed a consistent shift of the ratio between component motion percept duration and pattern motion percept duration in favor of component motion percepts (Figure 3E). These findings imply that the bulk of the effect that MECP2 duplication has on the perceptual reversals occurs due to increased stability of the component motion “transparent” percepts and a resulting shift of the ratio between component-pattern motion percept duration in favor of the component (“transparent”) interpretation. Ultra-stable component motion percepts then may contribute to lower probability to observe a reversal.

**Figure 3:**
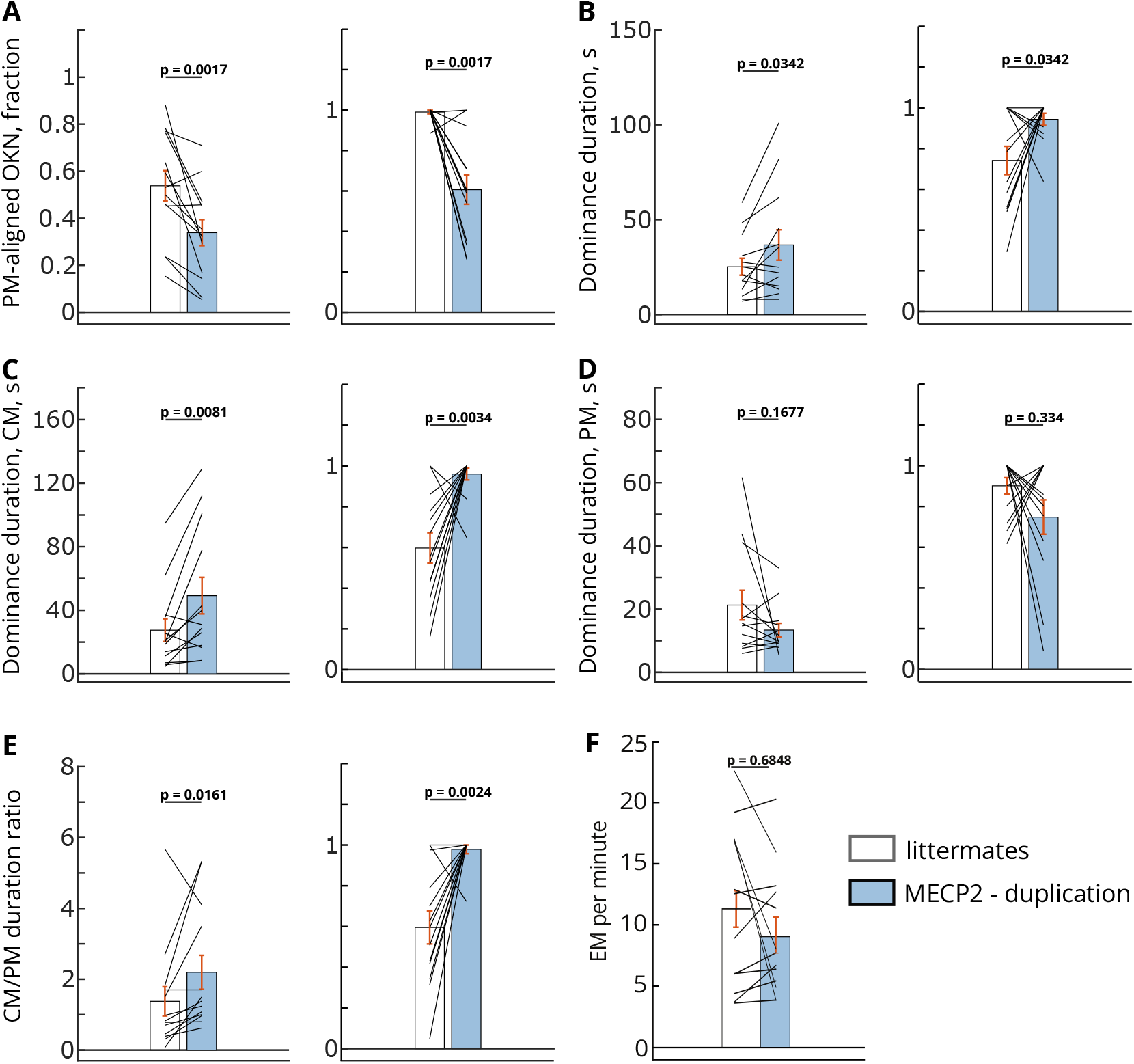
Atypical preference for local motion processing in MECP2 duplication syndrome. The reduced rate of perceptual reversals in MECP2 duplication mice is driven by the lengthening and over-stability of component motion (“transparent”) percepts. **A. The total fraction of nystagmoid eye movements aligned with pattern motion direction (“coherent” percept).** Even though there is considerable variance across data, in normal littermates (clear bar), nearly equal fractions of eye movements are aligned to either pattern motion direction (“coherent” percept, global motion) or component motion direction (“transparent” percept, local motion). In contrast, in MECP2 duplication mice, a greater portion of OKN eye movements is allocated to component local motion, and the fraction of pattern motion-aligned eye movements is reduced. Littermates, mean±sem: 0.538±0.064, median: 0.53; MECP2-ds, mean±sem: 0.339±0.055, median: 0.326. Left panel — raw data, right panel — data normalized by maximum inside each littermate — MECP2 duplication pair. **B. Dominance duration is increased in MECP2 duplication mice, following the decrease in reversal rate and reversal probability (Figure 2).** Littermates, mean ± sem: 25.2 ± 4.5, median: 21; MECP2-ds, mean ± sem: 36.7 ± 8, median: 25. Left panel — raw data, right panel — data normalized by maximum inside each littermate — MECP2 duplication pair; *p*-values, WSR. **C, D. The increase in average dominance duration is carried mainly by the increased durations of “transparent” percepts when the mouse is following the local motion of component gratings.** (**C**, littermates, mean ± sem: 27.5 ± 7.11, median: 20.2; MECP2-ds, mean ± sem: 49.2 ± 11.5, median: 31), while the global motion “coherent” percepts show inconsistent changes with shortening in some animals and lengthening in others (**D**, littermates, mean ± sem: 21.2 ± 4.7; MECP2-ds, mean ± sem: 13.36 ± 2.2). As a result, even though there is a general trend of shorter pattern-motion percepts in MECP2 duplication mice, it is not significant (*p* = 0.1677).

Figure 3: Left panels — raw data, right panels — data normalized by maximum inside each littermate — MECP2 duplication pair. **E. The ratio of dominance period durations is shifted in favor of transparent local motion percepts at the expense of global motion “coherent” percepts.** Littermates, mean ± sem: 1.376 ± 0.41, median: 0.82; MECP2-ds, mean ± sem: 2.2 ± 0.48, median: 1.38. Left panel — raw data, right panel — data normalized by maximum inside each littermate — MECP2 duplication pair. **F. The number of eye movements per minute in WT and MECP2-duplication mice.** Littermates, mean ± sem: 11.38 ± 1.81, median: 12.37; MECP2-ds, mean ± sem: 9,07 ±1.41, median: 7.54. These results indicate that the difference in frequency of eye movement is not significant (*p* = 0.6848). Left panel — raw data, right panel — data normalized by maximum inside each littermate — MECP2 duplication pair. White bars — littermates, blue bars — MECP2 duplication. All *p*-values are determined by two-sided Wilcoxon signed rank test (WSR), unless noted otherwise, *n* = 13 pairs.

## Discussion

### Slower dynamics of visual rivalry in MECP2 duplication syndrome

We view the world as generally stable even in the face of fast dynamic changes, such as fast-moving objects and emerging stimuli. This stability rests on an uncertain foundation: naturalistic scenes are inherently ambiguous, and the stable percepts of them are a result of a probabilistic process reflecting the most likely interpretation of the inputs. As a result, neuronal populations are constantly engaged in such ongoing interpretation and adjust their decision variables accordingly (***Aggelopoulos, 2015**; **Leopold and Logothetis, 1999**; **Sterzer et al., 2009***). In bi-stable and multi-stable perception, the competing interpretations of the sensory input cannot ultimately win against each other; as a result, the brain vacillates between the conflicting interpretations even though the stimulus stays the same. Visual rivalry involves a network of areas spanning V1, visual association areas, frontal lobe, supplementary motor cortex, and prefrontal cortex ***(Kleinschmidt et al., 1998**; **Knapen et al., 2011**; **Leopold and Logothetis, 1996**; **Lumer et al., 1998**;**Lumer and Rees, 1999)***. As a result, top-down cortical processes stemming from sensory-motor integration, attention and decision making affect the dynamics of visual rivalry. Perception, decision-making, and cognate sensory processing are pervasively impacted in neurological circuitopathies such as schizophrenia and autism (***Robertson et al., 2013**; **Heeger et al.,2017**; **Schmack et al., 2015***). Specifically, in idiopathic human autism, atypical sensory perception co-exists with higher-order deficits in social communication, cognitive flexibility, and executive function (***APA, 2013**;**Robertson and Baron-Cohen, 2017**; **Simmons et al., 2009**; **Van der Hallen et al., 2019***). As a distributed computation involving both the low-level sensation and perception processes and high-level processes pertaining on attention and decision-making, visual rivalry emerges as an attractive paradigm to study these processes and their interaction in the autism spectrum. In our study, we applied a monocular rivalry paradigm to explore if the dynamics of bistable visual perception were affected in the mouse model of MECP2 duplication syndrome of autism (***Collins et al., 2004***). This model reproduces some features of human autistic syndromes, including enhanced motor learning, motor, and visual stereotypies, and increased likelihood of seizure events (***Collins et al., 2004**; **Jiang et al., 2013**; **Samaco et al., 2012**; **Sztainberg et al., 2015**; **Zhang et al., 2017**; **Zhou et al., 2019**;**Ash et al., 2017**,**2021b**,a, **2022***). We found that the rate of perceptual reversals is decreased (Figure 2) in MECP2 duplication syndrome, while the average duration of individual percept dominance periods is prolonged (Figure 2). These effects occurred irrespective of the genetic line background of the mice, as we used both 129-MECP2 duplication line and FVB*C57 mixed background duplication line (***Ash et al., 2022***). Reduced rate of perceptual reversals under visual rivalry conditions in MECP2 duplication mice recapitulates the phenotype occurring in human idiopathic autism. ***(Robertson et al., 2013**; **Spiegel et al., 2019***). The magnitude of this reduction correlates with the expression of other autistic core traits, such as the severity of social communication deficits and ADOS score (***Spiegel et al., 2019***). Furthermore, in autistic subjects, slower binocular rivalry shares a common anatomical substrate with general cognitive rigidity — a part of the core repetitive restricted behaviors and interests (***Watanabe et al., 2019***).

### Atypical perception of visual motion in MECP2 duplication syndrome

Enhanced attention to visual detail and superior processing of local visual information are core traits of autism. Specifically, in autism, the visual perception is superior when the task is based on detecting local elements in the visual scene while the performance suffers when the subjects must focus on global elements (***Jolliffe and Baron-Cohen, 1997**; **Mottron et al., 1999**;**Happé et al., 2001**; **Plaisted et al., 1998**,**1999**; **Robertson and Baron-Cohen, 2017**; **Shah and Frith, 1983**;**Rinehart et al., 2000**; **Jarrold et al., 2005***). This perceptual phenotype is usually described in literature as “not seeing the forest behind the trees” (***Robertson et al., 2012**; **Frith, 2003***). Of particular relevance to our study are autism-related changes in the processing of visual motion and integration of local moving cues into a global moving percept (***Bertone et al., 2003**; **Brieber et al., 2010**;**Kaiser and Shiffrar, 2009**; **Koldewyn et al., 2010**; **Pellicano et al., 2005**; **Robertson et al., 2012**,**2014**;**Van der Hallen et al., 2019)***. The bi-stable-perception paradigm in our study makes use of two competing interpretations of a moving plaid: 1. the “transparent” interpretation where component gratings are seen as separate stimuli moving on top of each other, and 2. the “coherent” interpretation, where the stimulus is seen as a fusion of two moving component gratings resulting in a percept of moving pattern (***Adelson and Movshon, 1982**; **Castelo-Branco et al., 2000**; **Smith et al., 2005***). It is proposed that processing of complex stimuli like moving plaid rests on two distinct populations of neurons: orientation- and direction-selective component neurons and direction-of-motion selective pattern cells. While the first specialize on local motion information processing and responding to individual moving grating components, the latter ignore the orientation of the grating components, and instead respond to any stimulus moving in the preferred direction, including large-sized moving patterns such as naturalistic moving visual scenes. Pattern motion selectivity is posited to arise by integrating the inputs from component-motion-sensitive neurons. As one moves from primary visual areas to more specialized areas of the visual dorsal stream, the fraction of pattern cells and neurons integrating various types of local sensory information and computing global motion increases (***Albright and Stoner, 1995**; **Gizzi et al., 1990**; **Juavinett and Callaway, 2015**;**Khawaja et al., 2009**; **Movshon and Newsome, 1996**; **Palagina et al., 2017**; **Rust et al., 2006**; **Smith et al., 2005**; **Scannell et al., 1996**; **Rodman and Albright, 1989***). Pattern-motion processing in lower-ordervisual areas like V1 is strongly dependent on feedback from higher-order areas (***Guo et al., 2004***), while the integration of local motion cues into the global moving scenes by higher-order areas depends on the feedforward inputs from the V1 (***Movshon et al., 1985***).

Therefore, our competing interpretations are based on categorically different subtypes of visual motion: 1. local motion (when two individual component gratings are seen) and 2. global motion, occurring via integration of local motion cues and subsequent fusion of two gratings into a global moving pattern (as occurs in coherent moving plaid interpretation). Moreover, these two processes (global vs. local motion) are linked by feedforward and feedback connections across the cortical hierarchy.

In MECP2 duplication mice, we observed a pronounced preference for local motion percepts, both in terms of the fraction of eye movements aligned with component gratings and in terms of the duration of transparent versus coherent percepts (Figure 3). This recapitulates the visual motion processing peculiarities found in a subset of human subjects with autism ***(Robertson and Baron-Cohen, 2017**; **Van der Hallen et al., 2019***). Namely, studies using random dot kinematogram (RDK) display a subset of subjects with autism show increased motion coherence thresholds (e.g., a larger fraction of dots have to move together in the specified direction for the subject to detect coherent motion). However, this difference diminishes and disappears when the decision window is extended, implying that integration of local moving cues into a global moving percept is slowed down, but not fundamentally impaired or absent in autism spectrum (***Robertson et al., 2014***). Another group of studies found no differences in the behavioral performance of subjects when viewing RDK displays; however, subjects with autism still showed differential activation of visual areas in the dorsal stream, such as V1 and hMT, while observing and reporting coherent motion (***Brieber et al., 2010**; **Van der Hallen et al., 2019***). In a similar vein, our MECP2 duplication mice still consistently experience global moving pattern percepts. However, their durations show inconsistent changes: longer in one subgroup of MECP2 duplication animals and shorter in the others. While the duration of transparent percepts relying on local motion processing is consistently and dramatically increased compared to normal littermates (Figure 3).

### Interaction between the atypical perception of visual motion and reduced rate of perceptual reversals

In our paradigm, the MECP2 duplication mice show prolonged dominance periods of local motion perception. In contrast, the global motion percepts are generally shortened or unchanged, leading to shifted motion processing ratio favoring the local motion information over integrated motion information (Figure 3). Additionally, the total fraction of OKN eye movements aligned with component motion is greatly increased in MECP2 duplication, while the OKN fraction aligned with pattern motion is reduced (Figure 3). These observations imply that the capacity of neuronal populations reserved for the global motion percept formation and/or maintenance is reduced in MECP2 duplication syndrome, or the dynamics of such integration are altered. This is in line with two theories of autism — dorsal stream deficit theory (***Braddick et al., 2003**; **Greenaway et al., 2013**;**Macintyre-Beon et al., 2010**; **Chieffi, 2019***) and weak central coherence theory (***Dakin and Frith, 2005**;**Happé et al., 2001**; **Happé and Frith, 2006***). Dorsal stream deficit theory states that circuitry allocated to computing global motion from local moving cues is deficient in autism. In children with autism, this is exemplified by difficulties in following multiple moving objects simultaneously, impaired imitation of visual learning tasks, and performing complex movements without somatosensory feedback, since visual guidance of the motor output is disrupted (***Macintyre-Beon et al., 2010**;**Williams et al., 2004)***. Weak central coherence, on the other hand, proposes that global motion perception deficit may be due to a general cognitive style that prioritizes fine local details over global features (***Happé and Frith, 2006***). In both types of explanation, the preference of MECP2 duplication mice for local features atthe expense of globally coherent motion maybe a major contributor to diminished rate of visual rivalry. The bias for one specific rivaling interpretation of the stimulus may impair the ability of the brain to select an alternative interpretation and thus affect the rate of visual rivalry.

In MECP2 duplication, the coherent motion percepts appear to either not amass enough neuronal population activity or synchrony to remain stable, while local-motion percepts gain stability (Figure 3). Interestingly, the physiological basis for these changes may occur as early as primary visual cortical area V1 (***Ash et al., 2022**; **Palagina et al., 2017**; **Robertson et al., 2014***). First, pyramidal neurons in area V1 of MECP2 duplication mice show overly reliable firing in response to local motion information (for example, when moving gratings are used as a stimulus (***Ash et al., 2022***)). Second, area V1 harbors a significant portion of visual neurons dedicated to the processing of local motion and, in mice, contributes to the dynamics of bistable perception: removing V1 via lesion causes a decrease in the fraction of component motion-aligned OKN corresponding to local motion percepts (***Palagina et al., 2017***). In idiopathic human autism, hyperactivation of area V1 was found in a subset of subjects during the processing of coherent motion (***Brieber et al., 2010***). Additionally, in another subset of subjects with autism the areas of the dorsal stream, including V1 and middle temporal area, showed delayed activity during motion coherence processing ***(Robertson et al., 2014)***. Finally, neuronal responses of MECP2 duplication mice in area V1 show reduced coupling to ongoing cortical activity (***Ash et al., 2022***). This may result in disruption of both feedforward inputs from V1 to higher-order areas and weakening of the feedback from these higher-order areas to V1, reducing the integration of local motion cues there (***Ash et al., 2022***). Taken together, these observations point to an interesting possibility that the over-representation of local component motion in area V1 and disrupted connections between V1 and the rest of the visual dorsal stream are major contributors to the reduced rate of visual rivalry in autism. The reason is that they confer an advantage to the local motion information in the moving stimuli. In contrast, the synthesis of local information into the global motion of the scenes becomes impaired.

### Rate of visual rivalry, global motion synthesis and excitatory-inhibitory balance in cortical circuits

One of the prominent theories in autism states that core traits of the condition occur secondary to altered development of cortical interneurons and resulting shift in the balance between excitation and inhibition in cortical circuits across sensory and higher-order cortical areas ***(Casanova et al., 2003**; **Gogolla et al., 2009**; **Rubenstein and Merzenich, 2003**; **Robertson et al., 2014**,**2016***). Dynamics of visual rivalry and the rate of perceptual reversals are similarly hypothesized to depend on excitation-inhibition circuit wiring in the competing clusters of neurons coding for rivalrous percepts (***Laing and Chow, 2002**; **Seely and Chow, 2011**; **Klink et al., 2008a***). Computational models of binocular rivalry show that shifting excitatory-inhibitory ratio causes an increase in dominance durations, as eye-specific inputs maintain stable activity for more extended periods (***Dayan, 1998**; **Wilson, 2003**; **Klink et al., 2010**,**2008b**; **van Loon et al., 2013***). Altered local opponent inhibition in visuomotor areas was proposed to underlie the delayed integration of local moving features into global motion percepts in autism (***Robertson et al., 2014***). MECP2 dysfunction was shown to alter synchrony and net excitation-inhibition balance in neuronal circuits, with a greater impact on the phenotype of GABAergic interneurons. Overexpression of MECP2 was shown to affect predominantly genes affecting GABAergic signaling (***Cai et al., 2020**; **Chao et al., 2010***), with the result of disrupted synchronization within local and brain-wide networks (***Shou et al., 2017***). Thus, our findings that visual rivalry dynamics are slowed in MECP2 duplication mice and that they favor local motion percepts over global motion percepts are consistent with the altered excitation-inhibition dynamics theory of the autistic brain.

In summary, our MECP2 duplication mice phenotype reproduces core features of the autism spectrum — atypical perception of visual motion and slower dynamics of visual rivalry and thus can serve as a valid model of neural circuit dysfunction. Going forward, our bi-stable perception paradigm combined with 2-photon imaging and optogenetic manipulations (***Nikolenko et al., 2013**; **Sofroniew et al., 2016**; **Yizhar et al., 2011***) in the MECP2 duplication mouse model can be used to directly and causally test the following theories of the autism: excitatory-inhibitory imbalance, weak central coherence, dorsal stream deficiency and disrupted intracolumnar and cortex-wide connectivity.

## Acknowledgements

It is our pleasure to acknowledge Dr. Dmytro Bogatov for the valuable contributions to the computational part of this work. We are also grateful for Dr. Katerina Kalemaki, Dr. Andriani Rina, Lukas Braun, Dr. Joseph Lombardo, Dr. Mingyu Ye and Dr. Stefanos Chatzidakis and Dr. George Keliris for their advice and everyday support. This work was supported by NIH (Grant No: 1R21EY031537-01 and NS113890 - RO1).

## References

Adelson EH, Movshon JA. Phenomenal coherence of moving visual patterns. Nature. 1982 12; 300(5892):523–525. doi: 10.1038/300523a0.

Aggelopoulos NC. Perceptual inference. Neuroscience & Biobehavioral Reviews. 2015; 5:55–375. doi: 10.1016/j.neubiorev.2015.05.001.

Albright TD, Stoner GR. Visual motion perception. Proceedings of the National Academy of Sciences. 1995; 92(7):2433–2440. doi: 10.1073/pnas.92.7.2433.

APA. Diagnostic and Statistical Manual of mental disorders. 5 ed. American Psychiatric Publishing; 2013.

Ash RT, Palagina G, Fernandez-Leon JA, Park J, Seilheimer R, Lee S, Sabharwal J, Reyes F, Wang J, Lu D, Sarfraz M, Froudarakis E, Tolias AS, Wu SM, Smirnakis SM. Increased Reliability of Visually-Evoked Activity in Area V1 of the MECP2 duplication Mouse Model of Autism. Journal of Neuroscience. 2022; 42(33):6469–6482. doi: 10.1523/JNEUROSCI.0654-22.2022.

Ash RT, Buffington SA, Park J, Costa-Mattioli M, Zoghbi HY, Smirnakis SM. Excessive ERK-dependent synaptic clustering drives enhanced motor learning in the MECP2 duplication syndrome mouse model of autism. bioRxiv. 2017; doi: 10.1101/100875.

Ash RT, Buffington SA, Park J, Suter B, Costa-Mattioli M, Zoghbi HY, Smirnakis SM. Inhibition of Elevated Ras-MAPK Signaling Normalizes Enhanced Motor Learning and Excessive Clustered Dendritic Spine Stabilization in the MECP2 duplication Syndrome Mouse Model of Autism. eNeuro. 2021; 8(4).

Ash RT, Park J, Suter B, Zoghbi HY, Smirnakis SM. Excessive Formation and Stabilization of Dendritic Spine Clusters in the MECP2 duplication Syndrome Mouse Model of Autism. eNeuro. 2021; 8(1). doi: 10.1523/ENEURO.0282-20.2020.

Baron-Cohen S, Ashwin E, Ashwin C, Tavassoli T, Chakrabarti B. Talent in autism: hyper-systemizing, hyperattention to detail and sensory hyper-sensitivity. Philosophical Transactions of the Royal Society. 2009 5; 364(1522):1377–1383.

Bertone A, Mottron L, Jelenic P, Faubert J. Motion perception in autism: a “complex” issue. Journal of Cognitive Neuroscience. 2003 2; 15(2):218–225.

Bogatova D, Bogatov D, Palagina G, Dolia. Semi-automated time-series data processing Python 3 Software; 2023. https://github.com/mecp2-project/Dolia.

Bolton TAW, Morgenroth E, Preti MG, Van De Ville D. Tapping into Multi-Faceted Human Behavior and Psy-chopathology Using fMRI Brain Dynamics. Trends in Neurosciences. 2020 7; 43(9):667–680.

Braddick O, Atkinson J, Wattam-Bell J. Normal and anomalous development of visual motion processing: motion coherence and ‘dorsal-stream vulnerability,. Neuropsychologia. 2003; 41(13):1769–1784.

Brainard DH. The Psychophysics Toolbox. Spatial vision. 1997; 10(4):433–436.

Brieber S, Herpertz-Dahlmann B, Fink GR, Kamp-Becker I, Remschmidt H, Konrad K. Coherent motion processing in autism spectrum disorder (ASD): an fMRI study. Neuropsychologia. 2010 2; 48(6):1644–1651.

Cahill H, Nathans J. The optokinetic reflex as a tool for quantitative analyses of nervous system function in mice: application to genetic and drug-induced variation. PLoS One. 2008 4; 3(4):e2055.

Cai DC, Wang Z, Bo T, Yan S, Liu Y, Liu Z, Zeljic K, Chen X, Zhan Y, Xu X, Du Y, Wang Y, Cang J, Wang GZ, Zhang J, Sun Q, Qiu Z, Ge S, Ye Z, Wang Z. MECP2 duplication Causes Aberrant GABA Pathways, Circuits and Behaviors in Transgenic Monkeys: Neural Mappings to Patients with Autism. Journal of Neuroscience. 2020; 40(19):3799–3814. doi: 10.1523/JNEUROSCI.2727-19.2020.

Casanova MF, Buxhoeveden D, Gomez J. Disruption in the inhibitory architecture of the cell minicolumn: implications for autism. Neuroscientist. 2003 12; 9(6):496–507.

Castelo-Branco M, Goebel R, Neuenschwander S, Singer W. Neural synchrony correlates with surface segregation rules. Nature. 2000 6; 405(6787):685–689.

Chao HT, Chen H, Samaco RC, Xue M, Chahrour M, Yoo J, Neul JL, Gong S, Lu HC, Heintz N, Ekker M, Rubenstein JLR, Noebels JL, Rosenmund C, Zoghbi HY. Dysfunction in GABA signalling mediates autism-like stereotypies and Rett syndrome phenotypes. Nature. 2010 11; 468(7321):263–269.

Chieffi S. Dysfunction of Magnocellular/Dorsal Processing Stream in Schizophrenia. Current Psychiatry Reviews. 2019 01; 15. doi: 10.2174/1573400515666190119163522.

Collins AL, Levenson JM, Vilaythong AP, Richman R, Armstrong DL, Noebels JL, David Sweatt J, Zoghbi HY. Mild overexpression of MeCP2 causes a progressive neurological disorder in mice. Human Molecular Genetics. 2004 09; 13(21):2679–2689.

Dakin S, Frith U. Vagaries of visual perception in autism. Neuron. 2005 11; 48(3):497–507.

Dayan P. A hierarchical model of binocular rivalry. Neural Computation. 1998 7; 10(5):1119–1135.

Enoksson P. Binocular rivalry and monocular dominance studied with optokinetic nystagmus. Acta Ophthalmol (Copenh). 1963; 41:544–563.

Fox R, Todd S, Bettinger LA. Optokinetic nystagmus as an objective indicator of binocular rivalry. Vision Research. 1975 7; 15(7):849–853.

Frith U. Autism: Explaining the Enigma. 2 ed. Cognitive Development, London, England: Blackwell; 2003.

Gao E, DeAngelis GC, Burkhalter A. Parallel Input Channels to Mouse Primary Visual Cortex. Journal of Neuro-science. 2010; 30(17):5912–5926. doi: 10.1523/JNEUROSCI.6456-09.2010.

Gizzi MS, Katz E, Schumer RA, Movshon JA. Selectivity for orientation and direction of motion of single neurons in cat striate and extrastriate visual cortex. Journal of Neurophysiology. 1990 6; 63(6):1529–1543.

Gogolla N, Leblanc JJ, Quast KB, Südhof TC, Fagiolini M, Hensch TK. Common circuit defect of excitatory-inhibitory balance in mouse models of autism. Journal of Neurodevelopmental Disorders. 2009 6; 1(2):172–181.

Greenaway R, Davis G, Plaisted-Grant K. Marked selective impairment in autism on an index of magnocellular function. Neuropsychologia. 2013 3; 51(4):592–600.

Grzadzinski R, Huerta M, Lord C. DSM-5 and autism spectrum disorders (ASDs): an opportunityfor identifying ASD subtypes. Molecular Autism. 2013 5; 4(1):12.

Guo K, Benson PJ, Blakemore C. Pattern motion is present in V1 of awake but not anaesthetized monkeys. European Journal of Neuroscience. 2004 2; 19(4):1055–1066.

Van der Hallen R, Manning C, Evers K, Wagemans J. Global motion perception in autism spectrum disorder: A meta-analysis. Journal of Autism and Developmental Disorders. 2019 12; 49(12):4901–4918.

Happé F, Briskman J, Frith U. Exploring the cognitive phenotype of autism: weak”central coherence” in parents and siblings of children with autism: I. Experimental tests. Journal of Child Psychology and Psychiatry. 2001 3; 42(3):299–307.

Happé F, Frith U. The weak coherence account: detail-focused cognitive style in autism spectrum disorders. Journal of Autism and Developmental Disorders. 2006 1; 36(1):5–25.

Heeger DJ, Behrmann M, Dinstein I. Vision as a beachhead. Biological Psychiatry. 2017 5; 81(10):832–837.

Jarrold C, Gilchrist ID, Bender A. Embedded figures detection in autism and typical development: preliminary evidence of a double dissociation in relationships with visual search. Developmental Science. 2005 7; 8(4):344–351.

Jiang M, Ash RT, Baker SA, Suter B, Ferguson A, Park J, Rudy J,Torsky SP, Chao HT, Zoghbi HY, Smirnakis SM. Den-dritic arborization and spine dynamics are abnormal in the mouse model of MECP2 duplication syndrome. Journal of Neuroscience. 2013 12; 33(50):19518–19533.

Jolliffe T, Baron-Cohen S. Are people with autism and Asperger syndrome faster than normal on the Embedded Figures Test? Journal of Child Psychology and Psychiatry. 1997 7; 38(5):527–534.

Juavinett AL, Callaway EM. Pattern and component motion responses in mouse visual cortical areas. Current Biology. 2015 6; 25(13):1759–1764.

Kaiser MD, Shiffrar M. The visual perception of motion by observers with autism spectrum disorders: a review and synthesis. Psychonomic Bulletin & Review. 2009 10; 16(5):761–777.

Khawaja FA, Tsui JMG, Pack CC. Pattern motion selectivity of spiking outputs and local field potentials in macaque visual cortex. Journal of Neuroscience. 2009 10; 29(43):13702–13709.

Kleinschmidt A, Büchel C, Zeki S, Frackowiak RS. Human brain activity during spontaneously reversing perception of ambiguous figures. Proceedings of the Royal Society. 1998 12; 265(1413):2427–2433.

Klink PC, van Ee R, Nijs MM, Brouwer GJ, Noest AJ, van Wezel RJA. Early interactions between neuronal adaptation and voluntary control determine perceptual choices in bistable vision. Journal of Vision. 2008 05; 8(5):16–16. doi: 10.1167/8.5.16.

Klink PC, Brascamp JW, Blake R, van Wezel RJA. Experience-driven plasticity in binocular vision. Current Biology. 2010 8; 20(16):1464–1469.

Klink PC, van Ee R, van Wezel RJA. General Validity of Levelt,s Propositions Reveals Common Computational Mechanisms for Visual Rivalry. PLOS ONE. 2008 10; 3(10):1–9. doi: 10.1371/journal.pone.0003473.

Knapen T, Brascamp J, Pearson J, van Ee R, Blake R. The role of frontal and parietal brain areas in bistable perception. Journal of Neuroscience. 2011 7; 31(28):10293–10301.

Koldewyn K, Whitney D, Rivera SM. The psychophysics of visual motion and global form processing in autism. Brain. 2010 2; 133(Pt 2):599–610.

Laing CR, Chow CC. A spiking neuron model for binocular rivalry. Journal of Computational Neuroscience. 2002 1; 12(1):39–53.

Lawson RA, Papadakis AA, Higginson CI, Barnett JE, Wills MC, Strang JF, Wallace GL, Kenworthy L. Everyday executive function impairments predict comorbid psychopathology in autism spectrum and attention deficit hyperactivity disorders. Neuropsychology. 2015 5; 29(3):445–453.

Leopold DA, Logothetis NK. Activity changes in early visual cortex reflect monkeys, percepts during binocular rivalry. Nature. 1996 2; 379(6565):549–553.

Leopold DA, Logothetis NK. Multistable phenomena: changing views in perception. Trends in Cognitive Sciences. 1999 7; 3(7):254–264.

Leopold D, Fitzgibbons J, Logothetis N. The Role of Attention in Binocular Rivalry as Revealed through Optokinetic Nystagmus. Massachusetts Institute Of Technology Cambridge Artificial Intelligence Lab; 1995.

Logothetis N, Schall J. Neural correlates of subjective visual perception.. 1989 09; 245:761–3.

van Loon AM, Knapen T, Scholte HS, St John-Saaltink E, Donner TH, Lamme VAF. GABA shapes the dynamics of bistable perception. Current Biology. 2013 5; 23(9):823–827.

Lumer ED, Friston KJ, Rees G. Neural correlates of perceptual rivalry in the human brain. Science. 1998 6; 280(5371):1930–1934.

Lumer ED, Rees G. Covariation of activity in visual and prefrontal cortex associated with subjective visual perception. Proceedings of the National Academy of Sciences. 1999 2; 96(4):1669–1673.

Macintyre-Beon C, Ibrahim H, Hay I, Cockburn D, Calvert J, N Dutton G, Bowman R. Dorsal Stream Dysfunction in Children. A Review and an Approach to Diagnosis and Management. Current Pediatric Reviews. 2010; 6(3):166–182.

Mathis A, Mamidanna P, Cury KM, Abe T, Murthy VN, Mathis MW, Bethge M. DeepLabCut: markerless pose estimation of user-defined body parts with deep learning. Nature Neuroscience. 2018 9; 21(9):1281–1289.

Moreno-Bote R, Shpiro A, Rinzel J, Rubin N. Alternation rate in perceptual bistability is maximal at and symmetric around equi-dominance. Journal of Vision. 2010 9; 10(11):1.

Mottron L, Burack JA, Stauder JE, Robaey P. Perceptual processing among high-functioning persons with autism. Journal of Child Psychology and Psychiatry. 1999 2; 40(2):203–211.

Movshon J, Adelson E, Gizzi M, Newsome W. In: Chagas C, Gattass R, Gross C, editors. The analysis of moving visual patterns Pontificiae Academiae Scientiarum Scripta Varia, Vatican Press; 1985. p. 117–151.

Movshon JA, Newsome WT. Visual response properties of striate cortical neurons projecting to area MT in macaque monkeys. Journal of Neuroscience. 1996 12; 16(23):7733–7741.

Naber M, Frässle S, Einhäuser W. Perceptual rivalry: reflexes reveal the gradual nature of visual awareness. PLoS One. 2011 6; 6(6):e20910.

Niell CM, Stryker MP. Highly selective receptive fields in mouse visual cortex. Journal of Neuroscience. 2008 7; 28(30):7520–7536.

Nikolenko V, Peterka DS, Araya R, Woodruff A, Yuste R. Spatial light modulator microscopy. Cold Spring Harbor Protocols. 2013 12; 2013(12):1132–1141.

Ohki K, Chung S, Ch'ng YH, Kara P, Reid RC. Functional imaging with cellular resolution reveals precise microarchitecture in visual cortex. Nature. 2005 2; 433(7026):597–603.

Palagina G, Meyer JF, Smirnakis SM. Complex visual motion representation in mouse area V1. Journal of Neuroscience. 2017 1; 37(1):164–183.

Pellicano E, Gibson L, Maybery M, Durkin K, Badcock DR. Abnormal global processing along the dorsal visual pathway in autism: a possible mechanism for weak visuospatial coherence? Neuropsychologia. 2005; 43(7):1044–1053.

Peters SU, Hundley RJ, Wilson AK, Warren Z, Vehorn A, Carvalho CMB, Lupski JR, Ramocki MB. The behavioral phenotype in MECP2 duplication syndrome: a comparison with idiopathic autism. Autism Research. 2013 2; 6(1):42–50.

Plaisted K, O’Riordan M, Baron-Cohen S. Enhanced discrimination of novel, highly similar stimuli by adults with autism during a perceptual learning task. Journal of Child Psychology and Psychiatry. 1998 7; 39(5):765–775.

Plaisted K, Swettenham J, Rees L. Children with autism show local precedence in a divided attention task and global precedence in a selective attention task. Journal of Child Psychology and Psychiatry. 1999 7; 40(5):733–742.

Ramocki MB, Tavyev YJ, Peters SU. The MECP2 duplication syndrome. American Journal of Medical Genetics-Part A. 2010 5; 152A(5):1079–1088.

Rinehart NJ, Bradshaw JL, Moss SA, Brereton AV, Tonge BJ. Atypical interference of local detail on global processing in high-functioning autism and Asperger,s disorder. J Child Psychol Psychiatry. 2000 Sep; 41(6):769–778.

Robertson AE, Simmons DR. The sensory experiences of adults with autism spectrum disorder: A qualitative analysis. Perception. 2015; 44(5):569–586.

Robertson CE, Baron-Cohen S. Sensory perception in autism. Nature Reviews Neuroscience. 2017 11; 18(11):671–684.

Robertson CE, Kravitz DJ, Freyberg J, Baron-Cohen S, Baker CI. Slower rate of binocular rivalry in autism. Journal of Neuroscience. 2013 10; 33(43):16983–16991.

Robertson CE, Martin A, Baker CI, Baron-Cohen S. Atypical integration of motion signals in Autism Spectrum Conditions. PLoS One. 2012 11; 7(11):e48173.

Robertson CE, Ratai EM, Kanwisher N. Reduced GABAergic action in the autistic brain. Current Biology. 2016 1; 26(1):80–85.

Robertson CE, Thomas C, Kravitz DJ, Wallace GL, Baron-Cohen S, Martin A, Baker CI. Global motion perception deficits in autism are reflected as early as primary visual cortex. Brain. 2014 9; 137(Pt 9):2588–2599.

Rodman HR, Albright TD. Single-unit analysis of pattern-motion selective properties in the middle temporal visual area (MT). Exp Brain Res. 1989; 75(1):53–64.

Rubenstein JLR, Merzenich MM. Model of autism: increased ratio of excitation/inhibition in key neural systems. Genes, Brain and Behavior. 2003 10; 2(5):255–267.

Rust NC, Mante V, Simoncelli EP, Movshon JA. How MT cells analyze the motion of visual patterns. Nature Neuroscience. 2006 11; 9(11):1421–1431.

Samaco RC, Mandel-Brehm C, McGraw CM, Shaw CA, McGill BE, Zoghbi HY. Crh and Oprm1 mediate anxiety-related behavior and social approach in a mouse model of MECP2 duplication syndrome. Nature Genetics. 2012 1;44(2):206–211.

Scannell JW, Sengpiel F, Tovée MJ, Benson PJ, Blakemore C, Young MP. Visual motion processing in the anterior ectosylvian sulcus of the cat. Journal of Neurophysiology. 1996 8; 76(2):895–907.

Schmack K, Schnack A, Priller J, Sterzer P. Perceptual instability in schizophrenia: Probing predictive coding accounts of delusions with ambiguous stimuli. Schizophrenia Research: Cognition. 2015 6; 2(2):72–77.

Seely J, Chow CC. Role of mutual inhibition in binocular rivalry. Journal of Neurophysiology. 2011 11; 106(5):2136–2150.

Shafritz KM, Dichter GS, Baranek GT, Belger A. The neural circuitry mediating shifts in behavioral response and cognitive set in autism. Biological Psychiatry. 2008 5; 63(10):974–980.

Shah A, Frith U. An islet of ability in autistic children: a research note. Journal of Child Psychology and Psychiatry. 1983 10; 24(4):613–620.

Shou G, Mosconi MW, Wang J, Ethridge LE, Sweeney JA, Ding L. Electrophysiological signatures of atypical intrinsic brain connectivity networks in autism. Journal of Neural Engineering. 2017 8; 14(4):046010.

Simmons DR, Robertson AE, McKay LS, Toal E, McAleer P. Pollick FE. Vision in autism spectrum disorders. Vision Researchearch. 2009 11; 49(22):2705–2739.

Smith MA, Majaj NJ, Movshon JA. Dynamics of motion signaling by neurons in macaque area MT. Nature Neuroscience. 2005 2; 8(2):220–228.

Sofroniew NJ, Flickinger D, King J, Svoboda K. A large field of view two-photon mesoscope with subcellular resolution for in vivo imaging. Elife. 2016 6; 5.

Spiegel A, Mentch J, Haskins AJ, Robertson CE. Slower binocular rivalry in the autistic brain. Current Biology. 2019 9; 29(17):2948–2953.e3.

Sterzer P, Kleinschmidt A, Rees G. The neural bases of multistable perception. Trends in Cognitive Sciences. 2009 7; 13(7):310–318.

Sztainberg Y, Chen HM, Swann JW, Hao S, Tang B, Wu Z, TangJ, Wan YW, Liu Z, Rigo F, Zoghbi HY. Reversal of phenotypes in MECP2 duplication mice using genetic rescue or antisense oligonucleotides. Nature. 2015 12; 528(7580):123–126.

Ta D, Downs J, Baynam G, Wilson A, Richmond P, Leonard H. A brief history of MECP2 duplication syndrome: 20-years of clinical understanding. Orphanet Journal of Rare Diseases. 2022 3; 17(1):131.

Uddin LQ. Brain mechanisms supporting flexible cognition and behavior in adolescents with autism spectrum disorder. Biological Psychiatry. 2021 1; 89(2):172–183.

deVries SEJ, Lecoq JA, Buice MA, Groblewski PA, Ocker GK, Oliver M, Feng D, Cain N, Ledochowitsch P, Millman D, Roll K, Garrett M, Keenan T, Kuan L, Mihalas S, Olsen S, Thompson C, Wakeman W, Waters J, Williams D, et al. A large-scale standardized physiological survey reveals functional organization of the mouse visual cortex. Nature Neuroscience. 2019 12; 23(1):138–151.

Watanabe K. Optokinetic nystagmus with spontaneous reversal of transparent motion perception. Experimental Brain Research. 1999 11; 129(1):156–160.

Watanabe T, Lawson RP, Walldén YSE, Rees G. A neuroanatomical substrate linking perceptual stability to cognitive rigidity in autism. Journal of Neuroscience. 2019 8; 39(33):6540–6554.

Wei M, Sun F. The alternation of optokinetic responses driven by moving stimuli in humans. Brain Research. 1998 12; 813(2):406–410.

Williams JHG, Whiten A, Singh T. A systematic review of action imitation in autistic spectrum disorder. Journal of Autism and Developmental Disorders. 2004 6; 34(3):285–299.

Wilson HR. Computational evidence for a rivalry hierarchy in vision. Proceedings of the National Academy of Sciences. 2003 11; 100(24):14499–14503.

Yizhar O, Fenno LE, Prigge M, Schneider F, Davidson TJ, O’Shea DJ, Sohal VS, Goshen I, Finkelstein J, Paz JT, Stehfest K, Fudim R, Ramakrishnan C, Huguenard JR, Hegemann P, Deisseroth K. Neocortical excita-tion/inhibition balance in information processing and social dysfunction. Nature. 2011 7; 477(7363):171–178.

Yo C, Demer JL. Two-dimensional optokinetic nystagmus induced by moving plaids and texture boundaries. Evidence for multiple visual pathways. Invest Ophthalmol Vis Sci. 1992 Jul; 33(8):2490–2500.

Zhang D, Yu B, Liu J, Jiang W, Xie T, Zhang R, Tong D, Qiu Z, Yao H. Altered visual cortical processing in a mouse model of MECP2 duplication syndrome. Scientific Reports. 2017 7; 7(1):6468.

Zhao H, Mao X, Zhu C, Zou X, Peng F, Yang W, Li B, Li G, Ge T, Cui R. GABAergic system dysfunction in autism spectrum disorders. Frontiers in Cell and Developmental Biology. 2021; 9:781327.

Zhou C, Yan S, Qian S, Wang Z, Shi Z, Xiong Y, Zhou Y. Atypical response properties of the auditory cortex of awake MECP2-overexpressing mice. Frontiers in Neuroscience. 2019 5; 13:439.

